# Diversity of receptor expression in central and peripheral mouse neurons estimated from single cell RNA sequencing

**DOI:** 10.1101/2021.01.22.427766

**Authors:** Andi Wangzhou, Candler Paige, Pradipta R. Ray, Gregory Dussor, Theodore J. Price

## Abstract

Because somatosensory PNS neurons, in particular nociceptors, are specially tuned to be able to detect a wide variety of both exogenous and endogenous signals, it is widely assumed that these neurons express a greater variety of receptor genes. Because cells detect such signals via cell surface receptors, we sought to formally test the hypothesis that PNS neurons might express a broader array of cell surface receptors than CNS neurons using existing single cell RNA sequencing resources from mouse. We focused our analysis on ion channels, G-protein coupled receptors (GPCRS), receptor tyrosine kinase and cytokine family receptors. In partial support of our hypothesis, we found that mouse PNS somatosensory, sympathetic and enteric neurons and CNS neurons have similar receptor expression diversity in families of receptors examined, with the exception of GPCRs and cytokine receptors which showed greater diversity in the PNS. Surprisingly, these differences were mostly driven by enteric and sympathetic neurons, not by somatosensory neurons or nociceptors. Secondary analysis revealed many receptors that are very specifically expressed in subsets of PNS neurons, including some that are unique among neurons for nociceptors. Finally, we sought to examine specific ligand-receptor interactions between T cells and PNS and CNS neurons. Again, we noted that most interactions between these cells are shared by CNS and PNS neurons despite the fact that T cells only enter the CNS under rare circumstances. Our findings demonstrate that both PNS and CNS neurons express an astonishing array of cell surface receptors and suggest that most neurons are tuned to receive signals from other cells types, in particular immune cells.

## Introduction

Neurons receive signals from other cells through soluble chemical signals that act on receptors expressed by the neuron. Via the signaling action of these receptors, neurons are able to convert a chemical signal into electrical impulses that are the basis for information spread within the nervous system. Somatosensory neurons in the peripheral nervous system (PNS) can theoretically respond to soluble signals from almost any cell type in the body. A subset of neurons in the dorsal root (DRG) and trigeminal ganglion (TG) that detect injurious or potentially harmful stimuli, called nociceptors, are thought to be particularly tuned to detecting signals from other cell types because these neurons are the body’s first defense against cellular damage, inflammation, and pathogens (Woolf and Ma, 2007; Dubin and Patapoutian, 2010; Thakur et al., 2014; Chiu et al., 2016). Neurons in the central nervous system (CNS) have a smaller number of cell types from which to detect soluble ligands, but whether or not this means that they express a smaller repertoire of receptors has not been examined in a systematic way. Recent evidence indicates that cortical neurons are able to detect soluble factors released from immune cells that infiltrate the brain’s meninges, with profound influences on behavior (Filiano et al., 2016; Da Mesquita et al., 2018; Alves de Lima et al., 2020). Moreover, CNS glia can take on phenotypes that are strikingly similar to peripheral immune cells (Khakh and Deneen, 2019; Prinz et al., 2019). This growing understanding of direct non-neuronal influences on neuronal activity within the CNS implies that the receptor diversity thought to be a key characteristic of PNS neurons may also be found in many CNS neurons.

RNA sequencing technologies, in particular single cell sequencing technologies, have fundamentally changed our understanding of cellular populations within tissues (Stark et al., 2019). We now have unbiased expression profiles of most of the cell types in the PNS and CNS, at least in the mouse, with very specific knowledge of gene markers for these cells (Usoskin et al., 2015; Tasic et al., 2018; Zeisel et al., 2018; Zheng et al., 2019; Sharma et al., 2020). This information is incredibly useful because it gives genetic access to cell populations through a variety of transgenic and viral-vector technologies. It also is allowing for a better understanding of cell type conservation across species to enhance translational and evolutionary studies. These datasets can be mined in many interesting ways to reach conclusions that were not part of the aim of the original analysis. To this end, we were surprised that we were unable to find any previous studies that specifically examined pan-transcriptomic receptor expression diversity between neurons in the PNS and CNS. An exception may be the olfactory system where these neurons are very well known to express an array of GPCRs that are specifically involved in olfaction (Buck and Axel, 1991; Julius and Nathans, 2012). The primary goal of our work described here was to experimentally test the assumption that PNS somatosensory neurons express a greater variety of receptors than other types of neurons, excluding olfactory neurons.

In the work described here, we sought to gain insight into the diversity of receptor expression in PNS and CNS neurons of the mouse using a variety of published single cell sequencing resources (Tabula Muris et al., 2018; Tasic et al., 2018; Zeisel et al., 2018). Our specific hypothesis was that PNS somatosensory, sympathetic and enteric neurons would express a wider array of ion channels, G-protein coupled receptors (GPCRs), tyrosine receptor kinases (TRKs) and cytokine receptors. In partial agreement with our expectation, we found that PNS neurons express a greater diversity of GPCRs and cytokine receptors than CNS neurons. However, both PNS and CNS neurons express a very broad array of receptors of all of these families and there were no differences in expression diversity within the TRK or ion channel classes for PNS or CNS neurons. A secondary outcome of our analysis is the discovery of subsets of receptors within each family that are very specifically expressed in subsets of PNS neurons in the mouse. Because these receptors are not detected in CNS neurons, they may represent a new set of mechanistic or therapeutic targets for diseases of the PNS. Overall, our findings point to the astonishing array of receptors that are expressed by neurons, a feature that is common to both CNS and PNS neurons.

## Materials and methods

### Datasets

#### 1) scRNA-seq dataset for CNS and PNS neurons

In order to minimize technical noise, we selected scRNA-seq generated by Zeisel et al. (Zeisel et al., 2018) where both mouse CNS and PNS tissues were collected and sequenced with the same methods in the same lab, and with similar sequencing depth across all cell-types. The cell types used include neurons and glial cells throughout the cortex and other brain regions including mid- and hind-brain structure. For the PNS, the datasets include neurons and other cell types from the DRG, the sympathetic ganglia and enteric nervous system. We did not include data from the olfactory system due to the well-known overrepresentation of GPCRs in those neurons. We used the expression values of individual cells from CNS and PNS, and their identified cell-types provided by the original publication (L5_All.loom) for our analysis.

#### 2) scRNA-seq dataset for CNS neurons

It is possible that, despite CNS neurons having a greater variety of receptors than PNS neurons, they have relative low expression level, and thus due to the low sequencing depth nature of scRNA-seq, less receptors are considered expressed in CNS than PNS. Here we selected a deeply sequenced scRNA-seq dataset of CNS neurons by Allen Institute (Tasic et al., 2018) to show the differences we observed between CNS and PNS neurons are not a technical artifact.

#### 3) scRNA-seq dataset for T-cells

With multiple T-cell scRNA-seq datasets available, we selected the Tabula Muris dataset (Tabula Muris et al., 2018). This is because the T-cells sequenced in this dataset were not selected by any specific experimental procedure and are T-cells resident to several specific tissues. T-cells were pooled from 4 tissues in mouse: fat, muscle, lung, and spleen. This dataset was used to create ligand-receptor interactomes between T-cells and different types of neurons, as described below.

### Trinarization score

Trinarization score was developed by Zeisel et al, the original authors of the study from which the scRNA-seq datasets of CNS and PNS neurons were sourced (Zeisel et al., 2018). The trinarization score is a posterior probability score that identifies whether a gene is detectable in a set of sequenced cells (typically belonging to the same or related cell types). Presence or absence of reads in each cell are modelled as Bernoulli trials, with a Beta prior. The integral of the conjugate beta posterior P(Θ > f) is calculated, where f is the fraction of cells in the subpopulation where the gene should be detected. Methodology details can be found in the Zeisel et al. paper. Here, we used f = 0.05 to identify if even a small subpopulation of a particular cell type has detected reads. Beta distribution parameters α = 1.5, β = 2 were used. Genes with the trinarization score > 0.95 were considered detected in the particular cell type. While we do not quantize gene expression in this analysis, the term trinarization score is used for consistency of nomenclature with respect to Zeisel *et al* (Zeisel et al., 2018).

### Receptor diversity compared between CNS and PNS neurons

We used lists of GPCR, ion channel, and TRK genes from previously published paper in our lab (Wangzhou et al., 2020b). The list of cytokine related receptor genes was generated by combining genes under these gene groups from HGNC database (Braschi et al., 2019): tumor necrosis factor receptor superfamily, interferon receptors, interleukin receptors, complement system. We selected a human gene ontology database to increase translational value for human studies. Thus, we used the mouse orthologs of the human gene list to create the lists used in this analysis.

### Identification of PNS enriched gene modules and the calculation of enrichment score

The scRNA-seq dataset from mousebrain.org (Zeisel et al., 2018) as used for this analysis. Cell-types with less than 4500 genes detected were excluded. Hierarchical clustering was performed using a correlation based distance metric (1 – Pearsons Correlation Coefficient) and average-linkage on genes based on their trinarization score across all cell-types identified in CNS and PNS. Gene modules enriched in PNS neurons were then identified through an enrichment score. Enrichment score was calculated by the ratio of the mean trinarization scores of one set of cell-types to another. eg. the enrichment score for PNS neurons vs. CNS neurons for a specific gene A was calculated as the following:

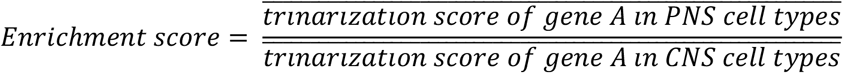

### Receptor diversity score

The diversity score is used to measure the diversity of genes under a certain gene class, GPCR, ion channel, TRK, or cytokine related receptors. The diversity score is calculated by summing the trinarization score of all genes under the corresponding gene class, for each cell-type. Greater diversity score is correlate with more genes are more likely to be expressed in the corresponding cell-type.

Trinarization scores, enrichment scores and diversity scores for genes of interest are available from Supplementary Files 1 – 5.

### Ligand-receptor interactions between T-cells and CNS and PNS neurons

We used scRNA-seq dataset from Tabula Muris (Tabula Muris et al., 2018) for the transcriptome of T-cells. Interactome between T-cells and CNS and PNS neurons were performed as previously described (Ray et al., 2020; Wangzhou et al., 2020a). Briefly, a ligand-receptor paired list was used to identify ligands expressed in T-cells. Then, we looked for the corresponding receptors of these ligands in CNS and PNS neurons. If the sum of trinarization score is greater than 25% of the cell-types, the gene is considered detected in CNS or PNS neurons. All identified interactions are listed in Supplementary File 5. We then further filtered these interactions for presentation in Figure 5C. We excluded all interactions where the receptor is considered detected in both CNS and PNS neurons. Then, for the PNS or CNS ‘specific’ receptors, we further looked at their corresponding ligands. If the ligands of these ‘specific’ receptors have other receptors that are considered detected in both CNS and PNS neurons, the ligand-receptor pair was excluded. The remaining was as presented in Figure 5C.

## Results

### Receptor diversity compared between CNS and PNS neurons for GPCR, ion channel, TRK and cytokine receptor families – mousebrain.org data

We first used the mousebrain.org dataset to explore receptor diversity between CNS and PNS neurons. We split CNS and PNS cell types into classifications described in (Zeisel et al., 2018) and mapped single cell expression by trinerization scores for all members of the GPCR family of receptors, excluding olfactory and other specialized receptor types that are mostly not expressed in either the CNS or DRG or enteric neurons. This revealed expression patterns for all GPCRs (Fig 1A, Supplementary File 1) across cell types in the CNS and PNS. Since a secondary goal of this analysis was to identify receptors that were exclusively expressed in the PNS in mouse, we performed hierarchical clustering on all GPCR genes by their expression across CNS and PNS neuron types, and identified this subcluster (green box in Fig 1A) and show these genes in more detail in Fig 1B, including enrichment scores for individual genes showing their relative degree of enrichment in the PNS. Some of these GPCR gene are well known to be enriched in sensory neurons, in particular the *Mrgpr* family (Dong et al., 2001; Zylka et al., 2003). Others, such as *F2r* and *F2rl2* (protease activated receptor type 1 and 2, respectively) have not been characterized as enriched for the PNS versus the CNS, but this receptor subfamily plays a well-established role in nociception (Vergnolle et al., 2001; Oikonomopoulou et al., 2018). Other sensory neuron enriched genes included *Lpar3* (Uchida et al., 2014; Velasco et al., 2017; Ueda, 2020) and P2ry2 (Moriyama et al., 2003; Stucky et al., 2004), both of which have also been implicated in nociception previously.

**Figure 1.**
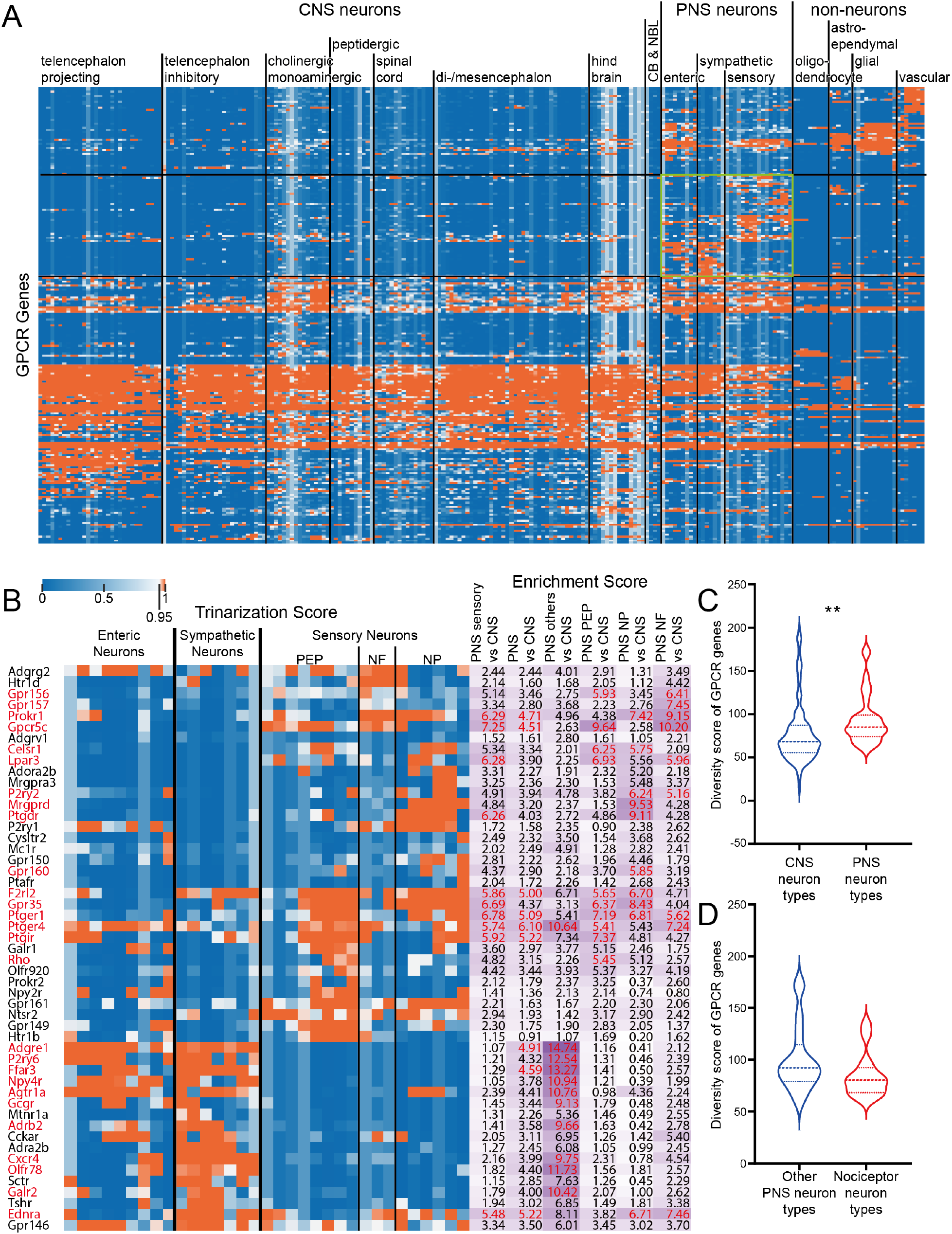
GPCR diversity between CNS and PNS neurons. A) Hierarchical clustering of all GPCR genes based on their trinarization score across all CNS and PNS cell-types. Green box highlights the GPCR genes enriched in PNS neurons. B) Detailed gene names and trinarization score across PNS neuron subtypes, and enrichment score for genes enriched in PNS neurons. Enrichment scores > 95^th^ percentile in the corresponding column are highlighted in red, along with the gene name. C) Violin plot showing the distribution of GPCR diversity scores for all CNS neuron types comparing with PNS neuron types. Welch’s t-test returns a p-value of 0.0073 indicating PNS neurons have greater GPCR diversity comparing with CNS neurons. D) Violin plot showing the distribution of GPCR diversity score for all non-nociceptor PNS neuron types comparing with nociceptors. Welch’s t-test returns a p-value of 0.0972.

Examining receptor diversity between CNS and PNS cell types, we found that overall, PNS neurons expressed more GPCRs than did CNS neurons (Fig 1C); however, it is notable that both cell types express a broad number of GPCRs and this diversity is consistent across most neuronal types examined in the Zeisel et al., dataset. When examining potential differences between nociceptor neuronal subtypes and all other PNS neurons profiled, we did not find any significant difference in receptor diversity (Fig 1D). A potential explanation for these findings is that all cell types, not just neurons, in the CNS and PNS express a large number of GPCRs. To test this, we utilized single cell sequencing for non-neuronal cell types in the Zeisel et al. dataset (Zeisel et al., 2018). Here we noted a dramatic difference in GPCR diversity between non-neuronal cells and PNS neurons where neurons expressed a far greater number of receptor genes (Supplementary Fig 1A).

We conducted a similar analysis for ion channel genes, including voltage gated-ion channels, even though many of them are technically not receptors. A great number of genes in this family were also broadly expressed across all neuron types (Fig 2A, Supplementary File 2), with a smaller subset of genes that were enriched in the PNS (Fig 2B). These genes included several voltage gated sodium channels which are well-known to be enriched in the DRG (e.g. *Scn10a (Akopian et al*., *1996)*), some purinergic ion channels like *P2rx2* and *P2rx3 (Bernier et al*., *2018; Burnstock, 2018)* and Trp channels that are also well-known sensory neuron-enriched genes (Basbaum et al., 2009). Unlike GPCRs, there were no significant differences between CNS and PNS neurons in ion channel expression diversity (Fig 2C), although there were PNS and CNS enriched genes, such as *Scn9a, Scn10a* and *Scn11a* in the PNS. There were also no significant differences in ion channel gene expression diversity between all other peripheral neuron types and nociceptors (Fig 2D). On the other hand, similar to GPCRs, there was a dramatic difference in ion channel diversity between PNS neurons and non-neuronal cell types with neurons expressing more ion channel genes (Supplementary Fig 1B).

**Figure 2.**
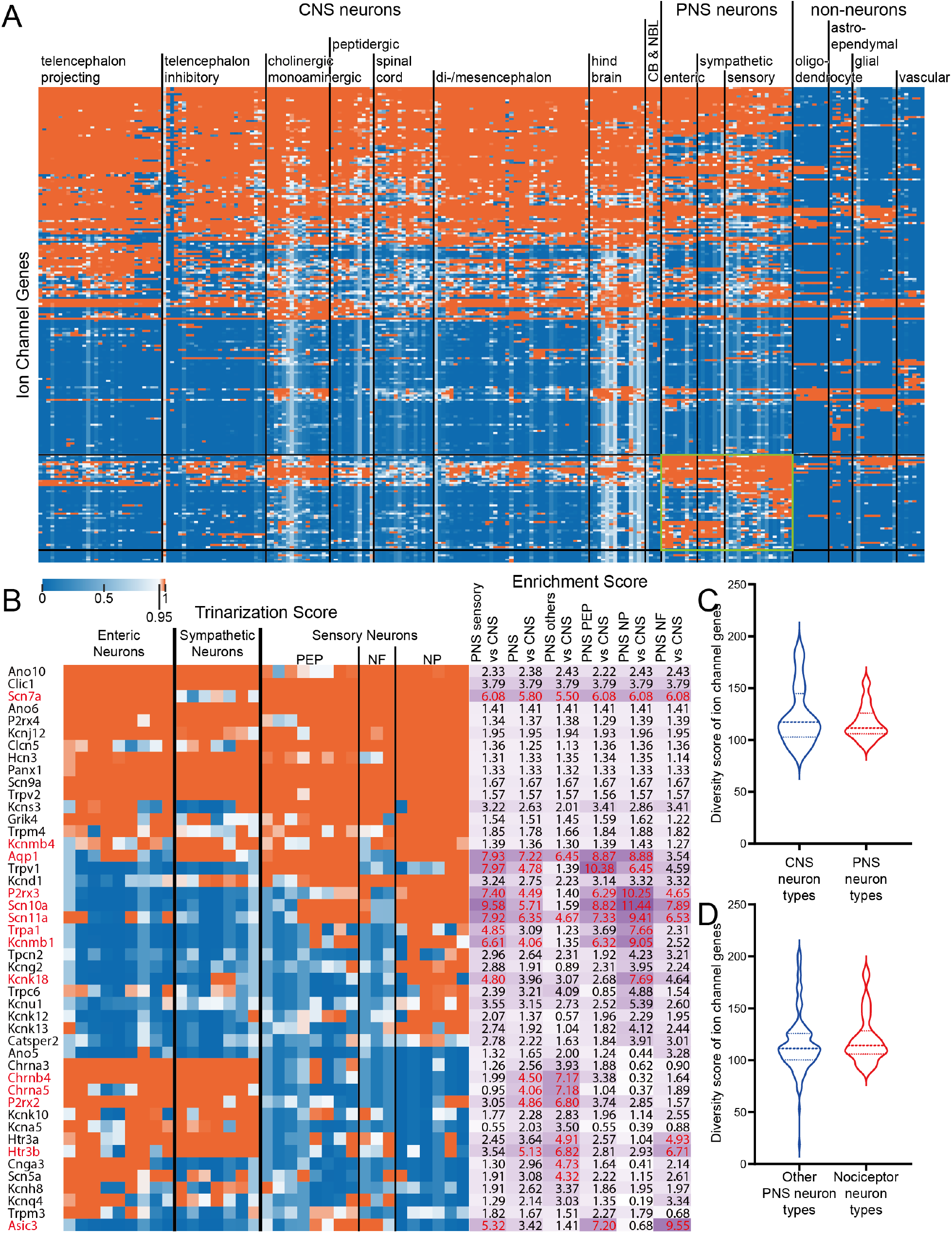
Ion channel diversity between CNS and PNS neurons. A) Hierarchical clustering of all ion channel genes based on their trinarization score across all CNS and PNS cell-types. Green box highlights the ion channel genes enriched in PNS neurons. B) Detailed gene names, trinarization score across PNS neuron subtypes, and enrichment scores for genes enriched in PNS neurons. Enrichment scores > 95^th^ percentile in the corresponding column are highlighted in red, along with the gene name. C) Violin plot showing the distribution of ion channel diversity score for all CNS neuron types comparing with PNS neuron types. Welch’s t-test returns a p-value of 0.3687. D) Violin plot showing the distribution of ion channel diversity score for all non-nociceptor PNS neuron types comparing with nociceptors. Welch’s t-test returns a p-value of 0.4115.

Among TRKs, we again noted broad expression across neuron subtypes for a good number of genes in this family (Fig 3A, Supplementary File 3), with a smaller subset of genes that were enriched in PNS neurons (Fig 3B). These TRK genes included the *Erbb3* gene which was exclusive to enteric neurons, where it plays a critical role in development (Espinosa-Medina et al., 2017), and the *Ntrk1* gene that was specific for sympathetic and sensory neuron clusters. The latter was expected given its genetic link to nociceptor and sympathetic neuron development (Lewin and Mendell, 1993; Lewin et al., 2014). There was not a difference in diversity of TRK expression between CNS and PNS neurons (Fig 3C) although we did note a decrease in diversity in nociceptors versus all other PNS neuron types (Fig 3D). Again, there was a substantially greater diversity of TRK expression in PNS neurons than in non-neuronal cell types (Supplementary Fig 1C).

**Figure 3.**
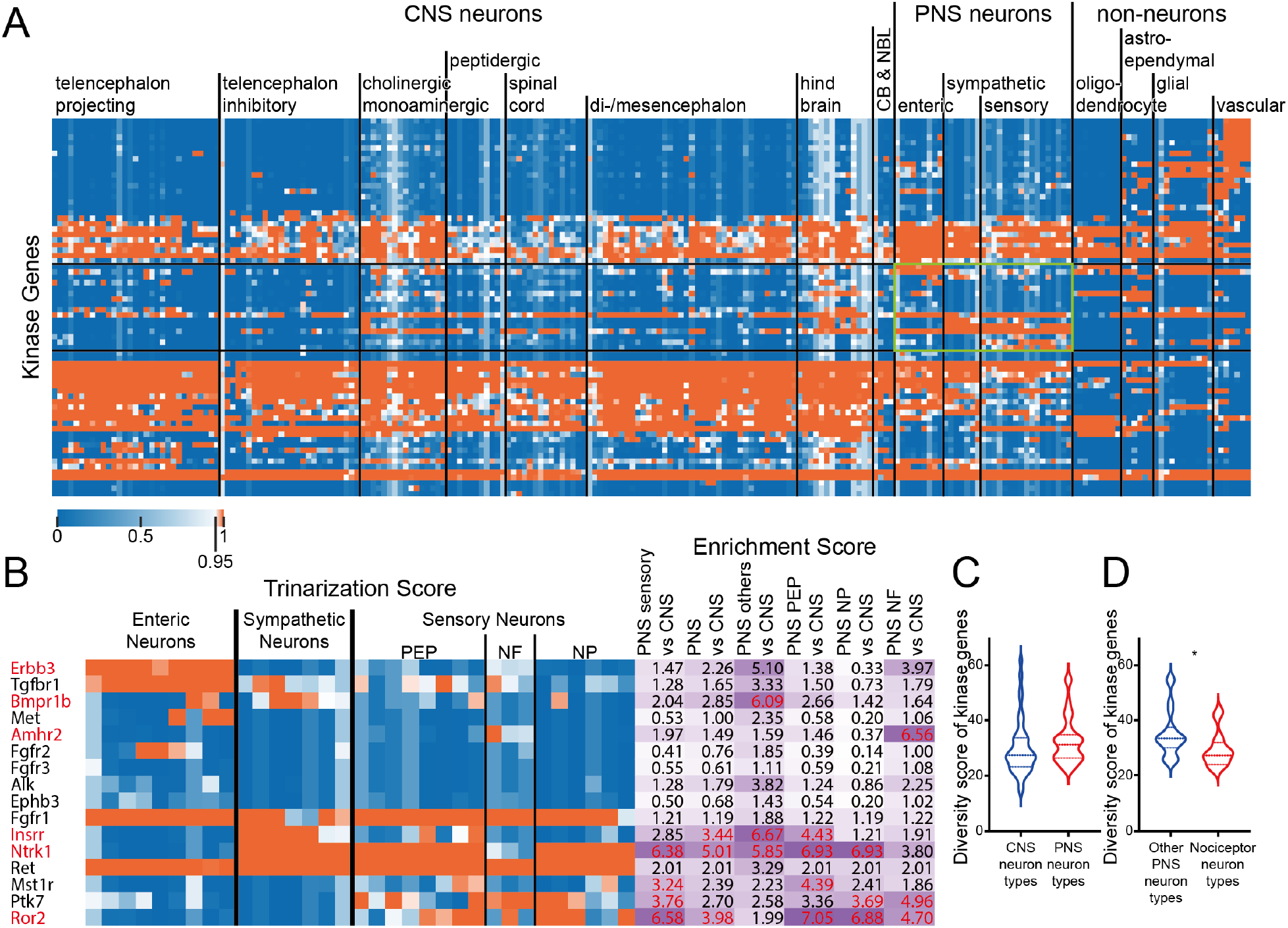
Tyrosine Receptor Kinase diversity between CNS and PNS neurons. A) Hierarchical clustering of all kinase genes based on their trinarization score across all CNS and PNS cell-types. Green box highlights the kinase genes enriched in PNS neurons. B) Detailed gene names, trinarization score across PNS neuron subtypes, and enrichment scores for genes enriched in PNS neurons. Enrichment scores > 95^th^ percentile in the corresponding column are highlighted in red, along with the gene name. C) Violin plot showing the distribution of kinase diversity score for all CNS neuron types comparing with PNS neuron types. Welch’s t-test returns a p-value of 0.1332. D) Violin plot showing the distribution of kinase diversity score for all non-nociceptor PNS neuron types comparing with nociceptors. Welch’s t-test returns a p-value of 0.0222.

Finally, we examined the cytokine receptor family (Fig 4A, Supplementary File 4). There was a clear population of receptors in this family that was enriched in PNS neurons (Fig 4B), including receptors for cytokines such as IL31, IL10 and interferons. This was reflected in a greater diversity in these receptors in PNS versus CNS neurons (Fig 4C), but this was mostly contributed by increased diversity in non-nociceptor PNS cell types (Fig 4D). Like all other receptor families, a greater diversity was noted in PNS neurons than in other non-neuronal cell types (Supplementary Figure 1D).

**Figure 4.**
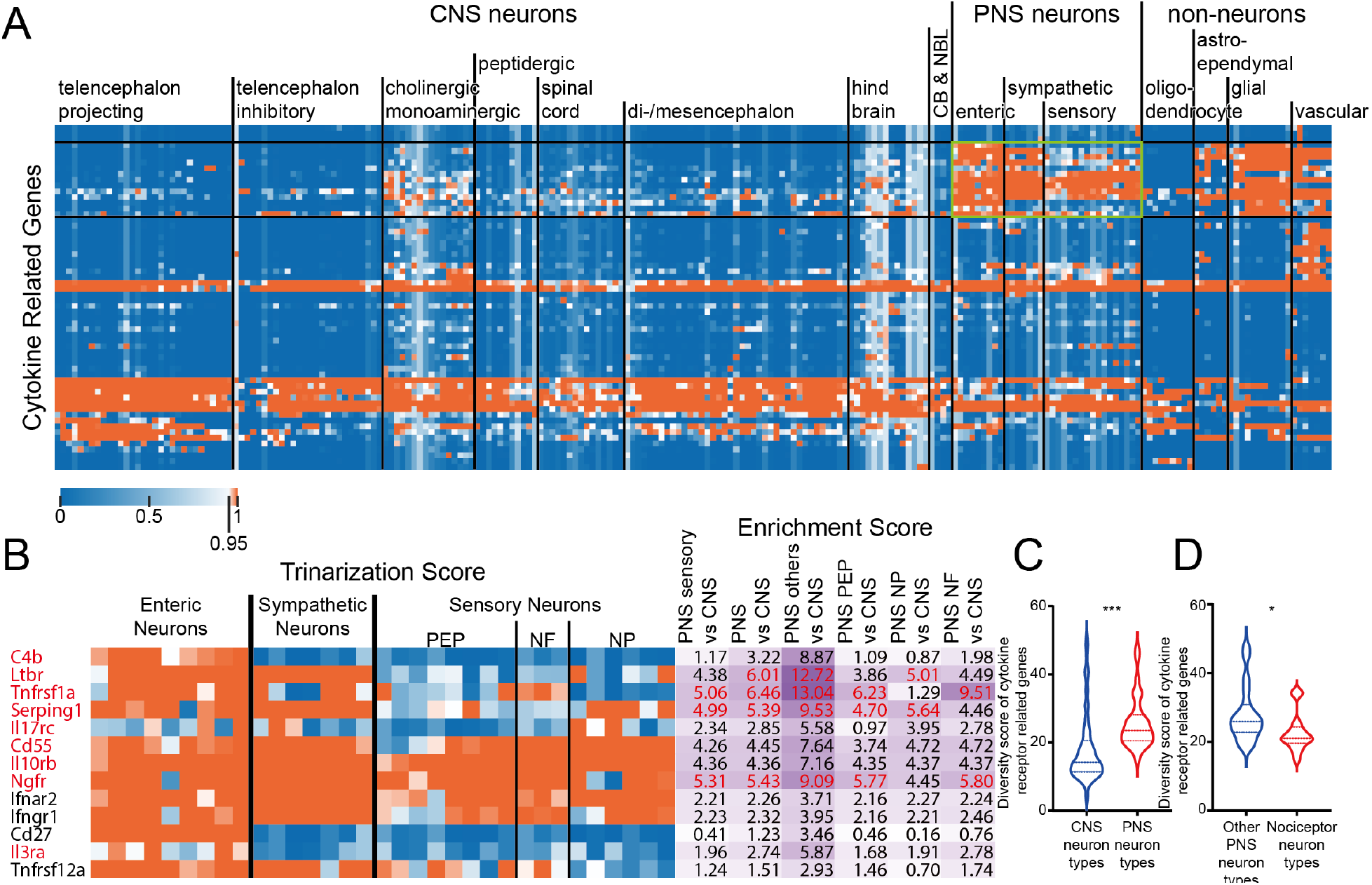
Cytokine related receptor diversity between CNS and PNS neurons. A) Hierarchical clustering of all cytokine related receptor genes based on their trinarization score across all CNS and PNS cell-types. Green box highlights the cytokine related receptor genes enriched in PNS neurons. B) Detailed gene names, trinarization score across PNS neuron subtypes, and enrichment scores for genes enriched in PNS neurons. Enrichment scores > 95^th^ percentile in the corresponding column are highlighted in red, along with the gene name. C) Violin plot showing the distribution of cytokine related receptor diversity score for all CNS neuron types comparing with PNS neuron types. Welch’s t-test returns a p-value of <0.0001 indicating that PNS neurons have greater cytokine related receptor diversity comparing with CNS neurons. D) Violin plot showing the distribution of cytokine related receptor diversity scores for all non-nociceptor PNS neuron types comparing with nociceptors. Welch’s t-test returns a p-value of 0.0335 indicating that there is less cytokine related receptor diversity detected in nociceptors than other PNS neurons.

**Figure 5.**
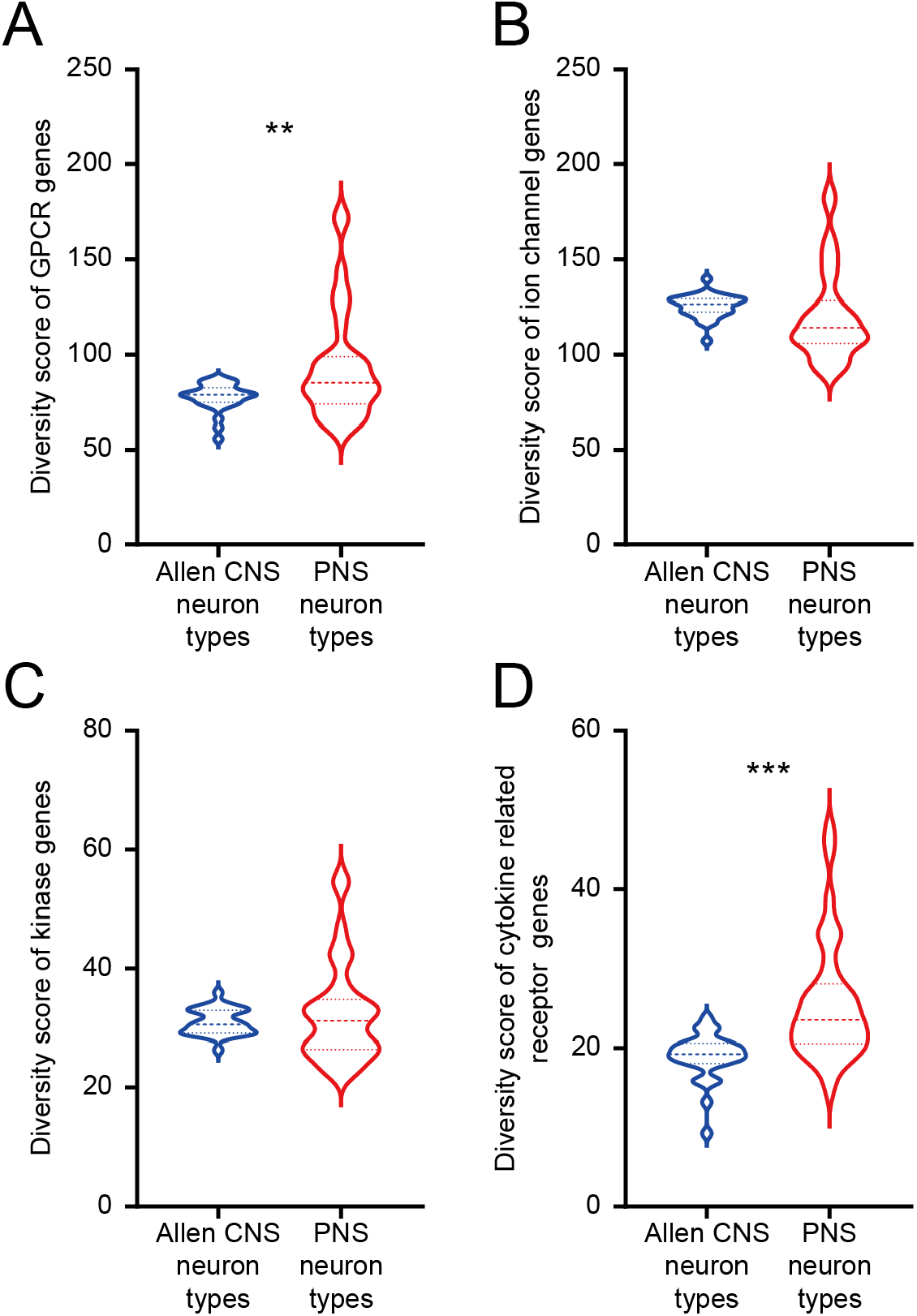
Gene diversity between CNS neurons from Allen brain dataset and PNS neurons from mousebrain.org dataset. A) Violin plot showing the distribution of GPCR diversity score for CNS neuron types and PNS neuron types. Welch’s t-test returns a p-value of 0.0039 indicating that PNS neurons have greater GPCR diversity compared with CNS neurons. B) Violin plot showing the distribution of ion channel diversity score for CNS neuron types and PNS neuron types. Welch’s t-test returns a p-value of 0.2289. C) Violin plot showing the distribution of tyrosine receptor kinase diversity scores for CNS neuron types and PNS neuron types. Welch’s t-test returns a p-value of 0.3131. D) Violin plot showing the distribution of cytokine related receptor diversity score for CNS neuron types and PNS neuron types. Welch’s t-test returns a p-value of <0.0001 indicating that PNS neurons have greater cytokine related receptor diversity compared with CNS neurons.

### Receptor diversity for PNS versus CNS cell types – Allen brain atlas data

A potential explanation for our findings is that this is an artifact of the CNS and PNS neuron preparations in the Zeisel et al. datasets (Zeisel et al., 2018). To formally test this possibility, we used single neuron sequencing data for CNS neurons from the Allen brain atlas dataset (Tasic et al., 2018). These neurons are sequenced more deeply than the Zeisel et al. dataset neurons so they could theoretically identify receptors in these families that are lowly expressed in neurons. Here we found precisely the same pattern that we found in the Zeisel et al. analysis. Despite the lower sequencing depth in Zeisel et al. dataset, PNS neurons still have greater diversity for GPCRs (Fig 5A) and cytokine recptor family genes (Fig 5D) than in the CNS whereas ion channels (Fig 5B) and TRKs were consistent (Fig 5C).

### Ligand-receptor interactions between T-cells and CNS and PNS neurons

Finally, we sought to understand if ligand-receptor interactions between a specific immune cell type and CNS and PNS neurons would be similar or different. We chose T-cells because they are mostly found outside of the nervous system but are increasingly recognized to play a critical role in many types of behavior. T-cell interaction with nociceptors is critical for both the development of pain and pain resolution (Sorge et al., 2015; Krukowski et al., 2016; Sommer et al., 2018; Laumet et al., 2019; Rosen et al., 2019; Kavelaars and Heijnen, 2021). T-cells in the meninges can have a profound impact on cortical neurons potentially promoting neurological disease (Filiano et al., 2016; Alves de Lima et al., 2020).

We used single cell transcriptomes of T-cells from different tissues in the tabula muris dataset (Tabula Muris et al., 2018). We used our previously described interactome framework (Wangzhou et al., 2020b; Wangzhou et al., 2020a) to identify ligands expressed in T-cells isolated from different mouse tissues. We found that most T-cells expressed a common set of ligands, with 84 ligands shared between muscle, fat, lung and spleen T-cells (Fig 6A). Based on these strong commonalities, we pooled all ligands found in these T-cells and intersected them with receptors (GPCRs, ion channels, TRKs and cytokine receptors) expressed either in PNS neurons or in cortical CNS neurons. We focused on cortical neurons because they are known to be influenced by T-cells in close proximity to these neurons in the meninges. There are other neurons, such as those that are found in the arcuate or subfornical areas, that do not have a blood brain barrier, but our goal was to focus on a broader group of cortical neurons rather than a specialized subset. In this ligand-receptor interactome, we identified 197 ligand receptor pairs (Fig 6B). Most of these pairs were shared by PNS and cortical neurons (107) and only 12 were unique to T-cells and cortical neurons (2 are shown in Fig 6C). Seventy-eight were unique to PNS neurons, and some of these are highlighted in Fig 6C and the entire interactome is shown in Supplementary File 5. Many of these PNS-specific interactions include receptors that are enriched in PNS neurons, such as *Ltbr* and *Tnfrsf1a*. Overall, these findings demonstrate that there are broad ligand-receptor interactions between both PNS neurons, which can come into direct contact with T-cells (Krukowski et al., 2016; Laumet et al., 2019), and CNS cortical neurons, which likely do not come into direct contact with T-cells, but rather communicate through release of factors in the meninges (Androdias et al., 2010; Filiano et al., 2016; Alves de Lima et al., 2020).

**Figure 6.**
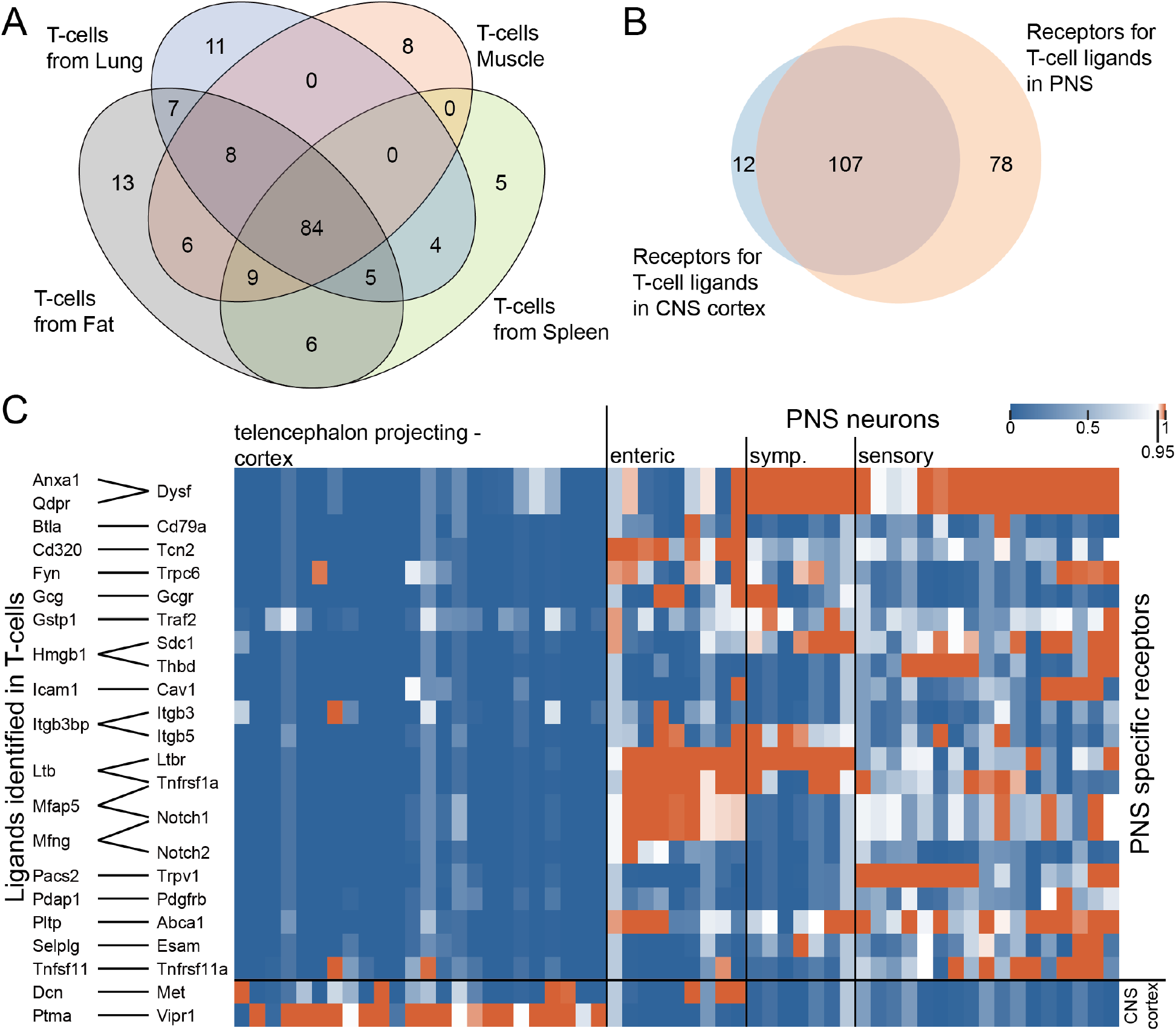
Ligand-receptor interactions identified between T-cells and CNS cortex or PNS neurons. A) Venn diagram showing the number of ligands identified in T-cells from fat, lung, muscle, and spleen, where corresponding receptors are detected in either CNS cortex or PNS neurons. B) Venn diagram showing the number of receptors identified in CNS cortex vs. PNS neurons where corresponding ligands are identified in T-cells. C) Ligand-receptor interactions where the receptor is specifically expressed either in CNS cortex or PNS neurons. Pairs with ligands having multiple receptor genes commonly expressed across CNS cortex and PNS neurons are not shown in the graph for clarity.

## Discussion

We set out to do these experiments with the hypothesis that peripheral neurons are likely to express a far greater diversity of receptors than CNS neurons. Our rationale for this hypothesis was simple, PNS neurons, in particular nociceptors, are able to respond to factors that can be released from almost any cell type in the body. Our findings show that PNS neurons, of all types, express an astonishing array of GPCRs, ion channels, TRKs and cytokine receptors. Surprisingly, CNS neurons showed similar diversity for most of these families of receptors. Even when there were significant differences in this diversity, such as GPCRs and cytokine receptors, this difference was small compared the difference between receptor diversity for PNS neurons and non-neuronal cell types. Therefore, while our findings provide some support for our original hypothesis, the weight of the evidence we collected in this study suggests that receptor expression diversity is similar between CNS and PNS neurons.

The next question is why would this be the case? There are several possible reasons. One is that the transcriptomic diversity of microglia, astrocytes and oligodendrocytes is far greater than was recognized prior to the emergence of single cell transcriptomic techniques (Khakh and Deneen, 2019; Prinz et al., 2019). These cells have long been recognized to participate in many aspects of disease, but we are also now learning about important roles that these cells play in normal physiology. An excellent recent example is the role of astrocytes in glutamate spillover in long-term potentiation (Henneberger et al., 2020). While the current evidence points mostly to glutamate clearance, it is likely that ligand-receptor interactions between neurons and astrocytes will play a key role in rapid structural changes in astrocytes around synapses. Another important new area of work is how immune cells that infiltrate the meninges can play a key role in shaping the activity of cortical neurons, and subsequently behavior. Currently, the best example of this is interferon gamma, which is secreted by meningeal T cells and then crosses the blood brain barrier, presumably via a transport mechanism, and acts on cortical neurons that express the receptor for this immune mediator (Filiano et al., 2016; Da Mesquita et al., 2018; Alves de Lima et al., 2020). Our work suggests that there are many such mediators from meningeal immune cells which could profoundly influence cortical neurons, if the factors can cross the blood brain barrier. Again, a primary conclusion of our work is that although there are some significant differences, PNS and CNS neurons are both well-tuned to respond to ligands that can be released from a large variety of cell types.

There are several limitations to our study. First, we have done this work using single cell sequencing resources from mouse. It will be important to do similar studies in human neurons, but single cell resources for the human PNS and/or CNS are not widely available. It will be interesting to make comparisons of receptor diversity across species. Based on previous work we have conducted looking at species differences in the DRG (Ray et al., 2018; Wangzhou et al., 2020b), we would expect that the diversity would be preserved, but that different specific receptors within families would likely be expressed between species. A second shortcoming is that we may have missed under-represented rare cell types in the PNS or, more likely, the CNS that may show dramatically different results than the cell types we have examined here. While these rare cell types would be unlikely to change our overall results, cell types that are potential outliers in their receptor diversity would be interesting to further analyze to understand the consequences of these differences. CNS neurons that are outside the blood brain barrier may be an interesting example of such outliers. A third limitation is that we have focused on datasets that represent healthy neurons. It is possible that receptor diversity could dramatically change in disease states. This will be a topic for future investigation. Our work creates a framework to do such an analysis. Extensive single cell sequencing datasets for injured peripheral neurons are now becoming available that will enable such future work (Hu et al., 2016; Nguyen et al., 2019; Renthal et al., 2020). A final limitation is that we have focused our analysis on receptor expression diversity within groups of neurons that have been classified by RNA sequencing, not on single neurons within any individual subset of cells. It would be interesting to approach this question of receptor diversity from the perspective of individual cells. However, current single cell sequencing technologies that are widely employed, such as nuclear RNA sequencing, likely do not provide a sufficiently robust snapshot of the transcriptome of single cells to do such an analysis (Stark et al., 2019). As these technologies continue to improve, these types of analyses can be done.

From the work described here, we reach the surprising conclusion that CNS and PNS neurons express similarly diverse repertoires of receptors, albeit with some exceptions depending on the receptor family. We suggest that most neurons are tuned to detect ligands expressed by a variety of cell types, a property that likely distinguishes them from many other cell types in the body. This does not mean that the expression diversity is identical in each type of neuron. In fact, our work identifies a large group of receptors that are exquisitely distinct for PNS neurons versus CNS neurons in the mouse. These receptors may represent a unique subset of drug targets for pain or other diseases if their distribution is conserved in humans.

## Notes

**Funding**: This work was supported by NIH grant NS065926 to TJP.

### Competing Interest Statement

AW, CP, PRR, GD and TJP are founders of Doloromics. The authors declare no other conflicts of interest.

